# Genome-wide CRISPRi screening reveals regulators of Alzheimer’s tau pathology shared between exosomal and vesicle-free tau seeds

**DOI:** 10.1101/2022.04.26.489622

**Authors:** Juan Carlos Polanco, Yevhen Akimov, Avinash Fernandes, Gabriel Rhys Hand, Adam Briner, Marloes van Roijen, Giuseppe Balistreri, Jürgen Götz

## Abstract

Aggregation of the microtubule-associated protein tau is a defining feature of Alzheimer’s disease and other tauopathies. Tau pathology is believed to be driven by both free tau aggregates and tau carried within exosomes, which propagate trans-synaptically and induce tau pathology in recipient neurons by a corrupting process of seeding. Here, we performed a genome-wide CRISPRi screen in tau biosensor cells and identified cellular regulators shared by both mechanisms of tau seeding. The top validated regulators are ANKLE2, BANF1, NUSAP1, EIF1AD, and VPS18, which work as factors that restrict tau aggregation initiated by both exosomal and vesicle-free tau seeds. Interestingly, ANKLE2 and BANF1 more robustly affected exosomal tau seeding than free aggregates. Lastly, validation studies revealed that several of the identified protein hits are downregulated in the brains of Alzheimer’s patients, suggesting that their decreased activity may be required for the emergence or progression of tau pathology in the human brain.

## Introduction

Tauopathies are neurodegenerative diseases in which the microtubule-associated protein tau undergoes a process of aggregation and fibrillization that gives rise to the pathological hallmark known as neurofibrillary tangles^1,2^. Alzheimer’s disease (AD) is a secondary tauopathy, with amyloid plaques featuring as an additional histopathological feature, which lacks in primary tauopathies such as frontotemporal lobar degeneration with tau (FTLD-tau), argyrophilic grain disease, progressive supranuclear palsy or corticobasal degeneration^1,2^. AD accounts for up to 80% of dementia cases worldwide, which currently affects more than 50 million people^2^. The pathological aggregation of tau impairs neuronal physiology at various levels, including axonal transport, action potential firing, synaptic plasticity, nuclear transport, chromatin structure, and mitochondrial function, which together lead to neurodegeneration and the ensuing cognitive and behavioral impairment^2-4^. A characteristic of tau aggregates, which are composed of misfolded tau modified by hyperphosphorylation and other post-translational modifications, is that they form proteopathic seeds that template the misfolding of physiological tau, and induce it to also misfold and form oligomers and fibrils^5,6^. This process is not restricted to the neurons in which the seeds form, but these can propagate trans-synaptically and be taken up by recipient neurons where they cause pathology by corrupting the native conformation of soluble tau^7^. This propagation requires release by donor cells and the subsequent internalization by recipient cells, which is brought about primarily via endocytosis. Importantly, to corrupt the conformation of physiological endogenous tau, the endocytosed tau seeds need to escape from the endolysosomes into the cytosol^8-10^.

Several studies have revealed that the propagation of tau seeds can be potentially controlled by interfering at multiple steps such as the production of the different tau seeds, their neuron-to-neuron transmission, internalization, endosomal escape into the cytosol, and cytoplasmic autophagy of newly-forming tau aggregates^7^. Fundamentally, two forms of tau seeds have been demonstrated to induce tau aggregation: (i) naked, i.e. vesicle-free tau in the form of oligomers or fibrils^11,12^, and (ii) tau encapsulated by the membranes of secretory extracellular vesicles known as exosomes^13-16^. There is an ongoing debate about which type of tau seed is critical in the progress of tau pathology. One recent report claimed using preparations from the same human AD brain tissue that exosomal tau seeds have higher transmissibility and cause more potent induction of tau pathology than vesicle-free tau seeds (whether oligomeric or fibrillar)^16^. However, any potential therapeutic approach that only targets either exosomes or free aggregates would confer incomplete protection from the tau pathology induced by the non-targeted seeds^7^. Therefore, it is crucial to identify the regulators of tau pathology that control both forms of tau seeding.

Here, we report novel regulators of seeded tau aggregation, which we discovered using a genome-wide CRISPR interference (CRISPRi) genetic screen^17^ coupled to a human cellular model of tau aggregation known as tau biosensor cells^11^ engineered to report tau aggregation by generating a FRET signal. Downstream bioinformatic analyses subsequently identified multiple hits, and we functionally validated some of our top hits, revealing that their individual knockdown predisposed the cells to tau aggregation. Interestingly, we found that several targets were also downregulated in the brains of AD patients suggesting that their decreased activity may be required for the emergence or progression of tau pathology in the human brain.

## Results

### Novel cellular regulators of tau pathology revealed by genome-wide CRISPRi screening

Tau biosensor cells^11^, designed to fluorescently display the extent of induced tau aggregation, have been widely used to study tau propagation and aggregation^10,18-20^. Moreover, we have shown previously that tau biosensor cells internalize both exosomes and vesicle-free tau isolated from brains of rTg4510 tau transgenic mice, resulting in the formation of intracellular tau inclusions^13^. To screen for novel regulators of tau pathology, we coupled tau biosensor cells to an optimized genome-wide CRISPRi library^17^ (Fig 1). First, we transduced these cells with a lentiviral KRAB-dCas9 construct, which expresses a nuclease-dead Cas9 (dCas9) fused to the transcription repressing KRAB domain to efficiently elicit gene silencing^17,21^. These modified tau biosensor cells (hereafter named BSKRAB cells) were then transduced with lentiviruses comprising the pooled whole-genome Dolcetto sgRNAs CRISPRi library^17^, and subsequently treated with sarkosyl-insoluble tau isolated from rTg4510 tau transgenic mouse brains^22^. We opted for this form of vesicle-free tau seeds which contains post-translational modifications such as phosphorylation (Fig 1a), given that previous studies have shown that sarkosyl-insoluble brain-derived aggregates are more potent seeders than aggregates generated *in vitro* with recombinant tau^23^. Indeed, 48 h after treatment of BSKRAB cells with sarkosyl-insoluble tau, we could detect robust seeded tau aggregation (Fig 1c). Importantly, all seeding assays were performed without lipofectamine, an agent that while strongly increasing seeded tau aggregation^13^, bypasses physiological vesicular trafficking by inducing and enhancing endolysosomal permeabilization^10^, which could create false-positive results in a genetic screen that uses tau aggregation as a readout.

**Fig 1:**
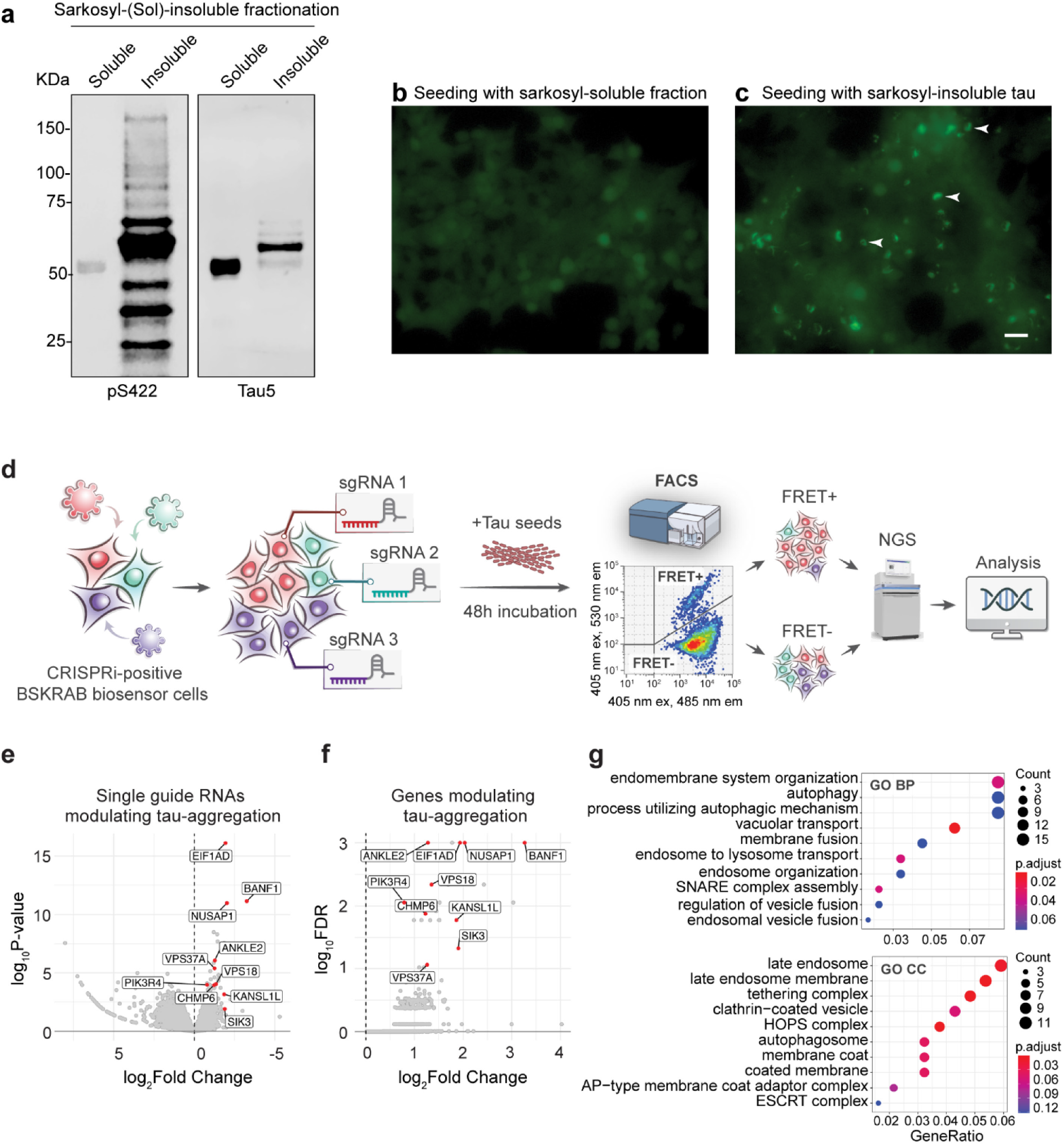
Unbiased discovery of regulators of tau aggregation using pooled CRISPRi screens and sarkosyl-insoluble tau. (**a**) Western blot analysis of sarkosyl-soluble and -insoluble fractions obtained from brains of P301L tau transgenic rTg4510 mice. The insoluble fraction exhibits phosphorylated tau (pS422) and high molecular weight total tau (Tau5). (**b-c**) Epifluorescence microscopy detecting tau RD-YFP in tau biosensor cells 48h after seeding with sarkosyl soluble (**b**) and insoluble tau (**c**). Brighter spots (arrowheads), representing tau aggregates appear only in cells that were treated with sarkosyl-insoluble tau. Scale bar: 50 μm. (**d**) Schematic representation of the pooled CRISPRi screen. Tau biosensor cells expressing the FRET pair tau RD-CFP and tau RD-YFP together with lentiviral KRAB-dCas9 (BSKRAB) were transduced with the Dolcetto CRISPRi library containing pooled lentiviral sgRNAs, targeting ∼18,000 genes, followed by incubation with sarkosyl-insoluble tau (vesicle-free tau seeds). 48 hours later, the cells were sorted into FRET(+) and FRET(–) populations using FACS. Samples were processed to generate an NGS sequencing library and sequenced on a NextSeq 500 instrument. (**e**) Volcano plot of the CRISPRi screen showing the enrichment of individual sgRNAs targeting distinct genes, highlighting in red some sgRNAs with high fold change. (**f**) Volcano plot showing gene-wise log2 fold changes of sgRNA counts versus false discovery rate (FDR). FDR values are based on robust ranking aggregation (RRA) from MAGeCK^24^. Genes that were followed up are highlighted in red. (**g**) Gene ontology (GO)-term enrichment analysis of the top 200 positive regulators of tau-aggregation identified by CRISPRi screening and enrichment for relevant pathways.

Using fluorescence-activated cell sorting (FACS) two cell populations were obtained, FRET-positive cells harboring induced tau aggregates and FRET-negative cells without aggregates. Genomic DNA was isolated, followed by next-generation sequencing (NGS) to quantify which sgRNAs were enriched in FRET-positive cells using the MAGeCK bioinformatic pipeline^24^ (Fig 1d). This revealed 23 genes (FDR<5%, Supplementary Table S1) that were positively enriched in FRET-positive cells with tau aggregation (Fig 1e,f). Furthermore, when the first 200 top gene hits, selected based on robust ranking aggregation (RRA) from MAGeCK^24^, were analyzed for pathway and protein complex enrichment using GO-BP (biological process) and GO-CC (cellular component) annotations, they were found to be primarily enriched in pathways of the late endosome, autophagosome, and tethering complexes coordinating endosome and lysosome fusion, suggesting an increase in tau aggregation due to a loss-of-function of genes with a role in autophagy and late endosomes (Fig 1g). By using a smaller CRISPRi library restricted to only genes within the endosomal pathway, two of our 23 top hits, CHMP6 and VPS13A (FDR<5%, Supplementary Table S1), have also been recently reported^8^. In contrast to that study, we used a genome-wide library enabling the identification of many additional hits.

### Validation of CRISPRi hits reveals genes that predispose cells to both vesicle-free and exosomal-tau seeded aggregation

We next functionally validated ten of our top hits (Fig 1f; ANKLE2, NUSAP1, EIF1AD, BANF1, VPS18, CHMP6, PIK3R4, KANSL1L, SIK3, and VPS37A) which were selected considering that they have been shown to be expressed in neurons of the human brain. Each gene was individually targeted with three sgRNAs from the Dolcetto library to achieve CRISPRi-mediated gene silencing in tau biosensor cells (hereafter named BSKRAB-KD), using a lentiviral vector in which both KRAB-dCas9 and the individual sgRNA were contained within a single construct. BSKRAB-KD cells in which individual genes had been silenced were then compared with a control that was obtained using the mean of three non-targeting sgRNAs (Fig 2a). For validation, the cells were treated with either vesicle-free tau seeds (sarkosyl-insoluble tau) or exosomal tau seeds isolated from rTg4510 mouse brains, followed by detection and quantification of tau aggregation using FRET flow cytometry (Fig 2a). We used one of the ten hits, CHMP6, for benchmarking purposes (Fig 2b), given that its knockdown has previously been reported to induce tau aggregation^8^. We found that apart from the CHMP6 knockdown (Fig 2b), the individual knockdowns of ANKLE2, BANF1, NUSAP1, EIF1AD as well as VPS18 strongly and significantly promoted tau aggregation induced by both exosomal and vesicle-free tau seeds (Fig 2c-g). Remarkably, ANKLE2 and BANF1 showed a substantially stronger effect on tau aggregation induced by exosomes (Fig 2c,d). Of note, silencing of ANKLE2 resulted in a remarkable forty-fold induction of seeding when treated with exosomes, compared to a six-fold induction with vesicle-free tau (Fig 2c).

**Fig 2:**
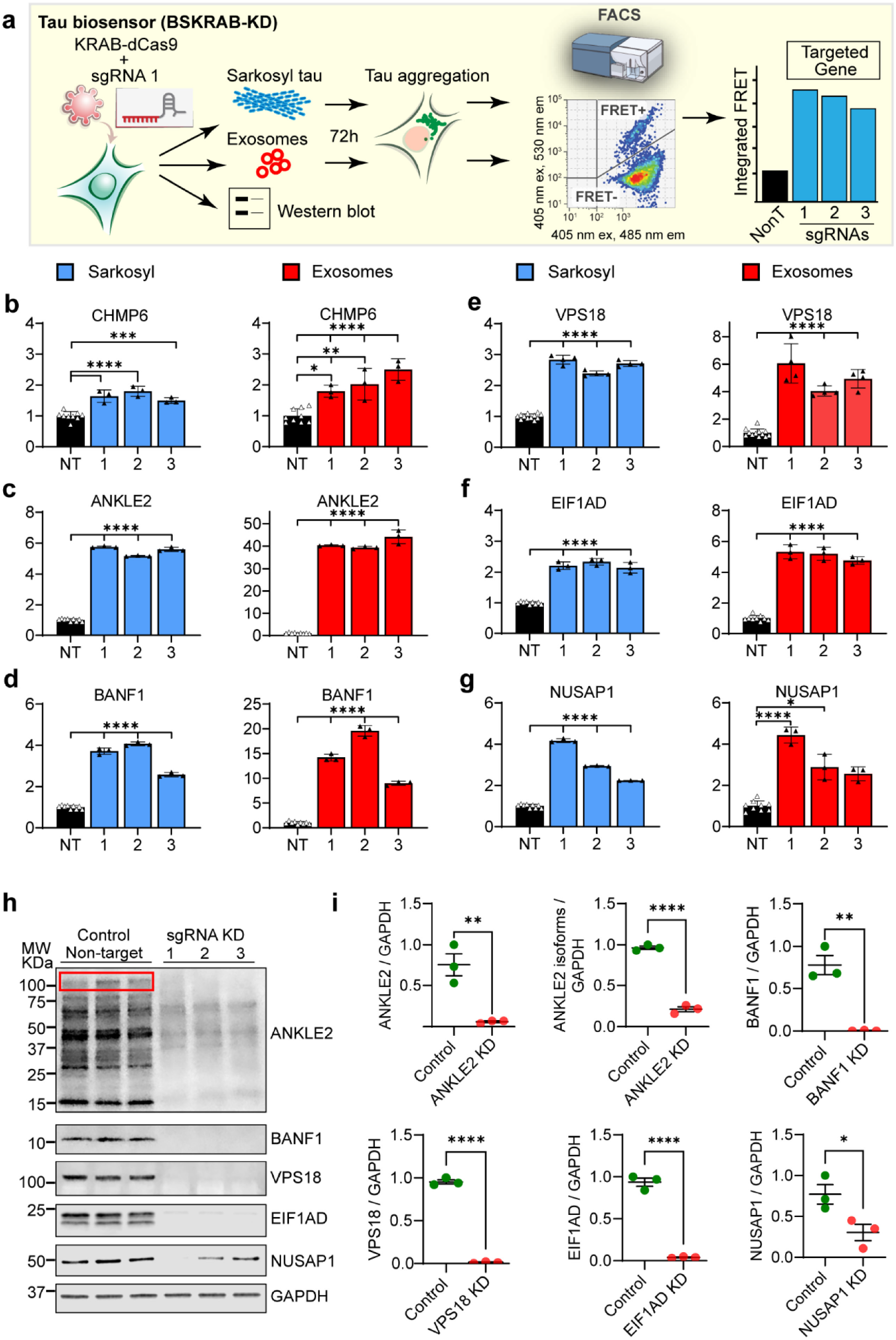
Functional validation of bioinformatic hits using individual CRISPRi knockdowns followed by incubation with exosomal and vesicle-free tau seeds. (**a**) Schematic representation of the workflow for functional validation. Individual sgRNAs were used to silence the corresponding genes in combination with KRAB-dCas9 in tau biosensor cells (BSKRAB-KD), then treated with either sarkosyl-insoluble tau (vesicle-free tau seeds) or exosomal tau seeds for 72 hours, followed by detection and quantification of tau aggregation using FRET flow cytometry. A fraction of the same BSKRAB-KD cells was also grown for 72 hours to corroborate knockdown of protein expression using western blots. (**b-g**) Integrated FRET intensities represent levels of tau aggregation upon knocking down the different targets. The control black column (NT) is the average obtained with three independent non-targeting sgRNAs (n=3) assessed in triplicates. Control cells were compared with knockdown cells targeted individually (1, 2, and 3). Error bars represent SEM for n=3, \**p<*0.05; \*\**p<*0.01; \*\*\**p<*0.001; \*\*\*\**p<*0.0001. Each single targeting sgRNA increased tau aggregation with both exosomal and vesicle-free tau seeds. (**c**,**d**) Interestingly, ANKLE2 and BANF1 appear to induce a stronger effect on tau aggregation induced by exosomes. (**h**) Quantitative western blot analysis of BSKRAB-KD knockdown cells. Each sgRNAs generated a protein knockdown of the targeted gene. Note that the ANKLE2-specific antibody reacts with several isoforms, including the canonical variant sized 104-117 kDa (red box outline), however, all isoforms were downregulated when the ANKLE2 locus was silenced. Similarly, the EIF1AD antibody recognized the canonical isoform of 19 kDa and one additional variant of lower molecular weight, both being silenced with the individual sgRNAs against EIF1AD. (**i**) Quantification of the extent of protein knockdown for the different targeted genes. Error bars represent SEM for n=3, *p<0.05; **p<0.01,, ****p<0.0001.

Furthermore, when whole-cell lysates from the five types of knockdown cells were analyzed by quantitative western blotting (Fig 2h), we found that each of the sgRNAs was able to induce gene silencing of the targeted genes (Fig 2i), demonstrating the reproducibility and robustness of CRISPRi-mediated gene silencing. Of note, we found that the polyclonal ANKLE2 antibody reacted with the canonical isoform of 104-117 KDa (Fig 2h, red box outline) and other isoforms of lower molecular weight, which were also downregulated when the ANKLE2 locus was silenced (Fig 2h). We also evaluated PIK3R4, KANSL1L and SIK3, but these hits did not potentiate tau aggregation in the validation assays (Supplementary Fig S1) and are likely false-positives. VPS37A exhibited a low potentiation and was therefore not followed up further (Supplementary Fig S1).

### Increased tau aggregation is not linked to mechanisms that affect tau uptake

Having demonstrated that the individual gene knockdowns of ANKLE2, BANF1, NUSAP1, EIF1AD, and VPS18 resulted in enhanced seeded tau aggregation (Fig 2), this raised the question whether such an increased tau aggregation could be merely caused by higher uptake of tau seeds. Therefore, to assess the level of internalization of tau seeds, sarkosyl-insoluble tau was labeled with the far-red dye Alexa-Fluor-647, whereas the membranes of exosomal tau seeds were labeled with the far-red fluorescent membrane probe CellVue Claret (CVC). Then, the level of uptake was measured in individual gene knockdowns of BSKRAB-KD cells using flow cytometry (Fig 3a). We found that none of the gene knockdowns impacted cellular uptake as shown for vesicle-free tau (Fig 3b) and exosomes (Fig 3c), indicating that the observed increases in tau aggregation (Fig 2) were not because of an increased internalization but rather a cell-autonomous mechanism downstream of the seed uptake.

**Fig 3:**
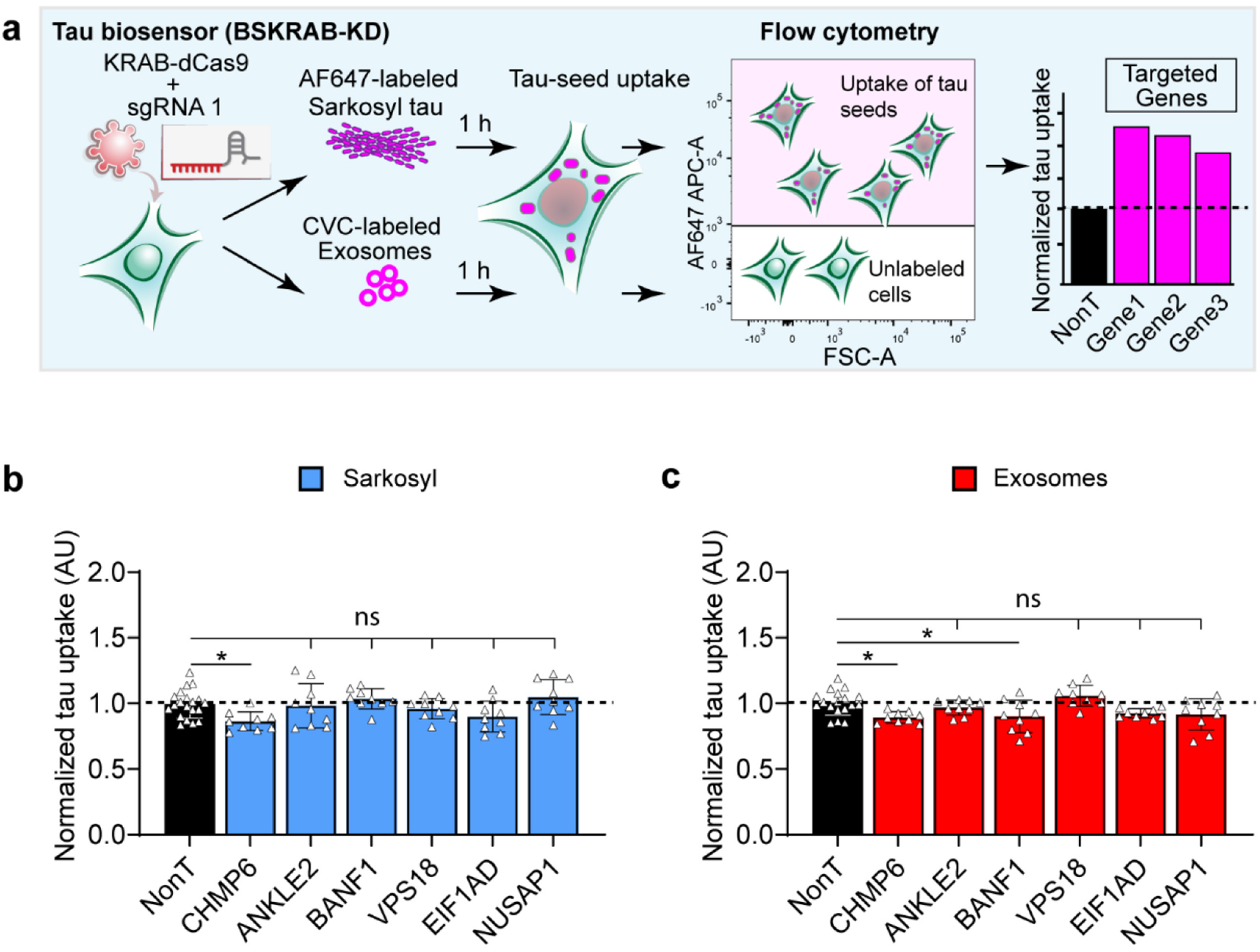
Analysis of the levels of tau-seed uptake for the different knockdown cells. (**a**) Schematic of the workflow for quantifying the uptake of exosomal and vesicle-free tau seeds. BSKRAB-KD knockdown cells were treated with far-red labeled tau seeds either AF647-labeled sarkosyl tau or CVC-labeled exosomes for 1 hour, followed by detection of cells harboring the labeled tau seeds using flow cytometry. Tau uptake was quantified by measuring the mean fluorescence intensity in far-red positive cells. (**b-c**) None of the gene knockdowns generated a significant increase in tau seed uptake (dashed line), suggesting that these genes are downstream of tau seed internalization, and therefore impact tau aggregation through cell-autonomous mechanisms. Three independent non-targeting sgRNAs (n=3) assayed in triplicates were used as control (black column) and compared with three knockdown sgRNAs targeting each gene, which were pooled for comparison. Error bars represent SEM for n= 9, *p<0.05; ns, non-significant.

### Decreased levels of VPS18, NUSAP1, and EIF1AD in the brains of Alzheimer’s patients

Validation studies in tau biosensor cells revealed that decreasing the levels of key cellular factors results in increased tau aggregation. To validate our hits further *in vivo*, we asked whether tau aggregation in AD patients correlates with the downregulation of the discovered and functionally validated CRISPRi targets. To address this, we performed a quantitative western blot analysis of *post mortem* brain samples from AD patients (Fig 4 and Table 1). Strengthening the relevance of our screen for the human condition, our analysis revealed that VPS18, NUSAP1, and EIF1AD were all downregulated in cortical AD brain tissue (Fig 4a, c-e), which is characterized by a substantial accumulation of phosphorylated and aggregated tau (Fig 4 b,g). Surprisingly, no statistically significant differences were found for ANKLE2 levels between AD and control patients (Fig 4f), although the data was heterogenous for the patients analyzed. Of note, only the predominant, canonical ANKLE2 104-117 kDa isoform (Fig 4a, red box outline) was quantified (Fig 4f). We did not detect BANF1 via western blotting in our cortical brain samples, possibly reflecting low expression levels. Together, our data suggest that the decreased activity of VPS18, NUSAP1, and EIF1AD in AD may be an upstream pathogenic event that enhances the aggregation and interneuronal spreading of pathological tau through the human brain.

**Table 1:**
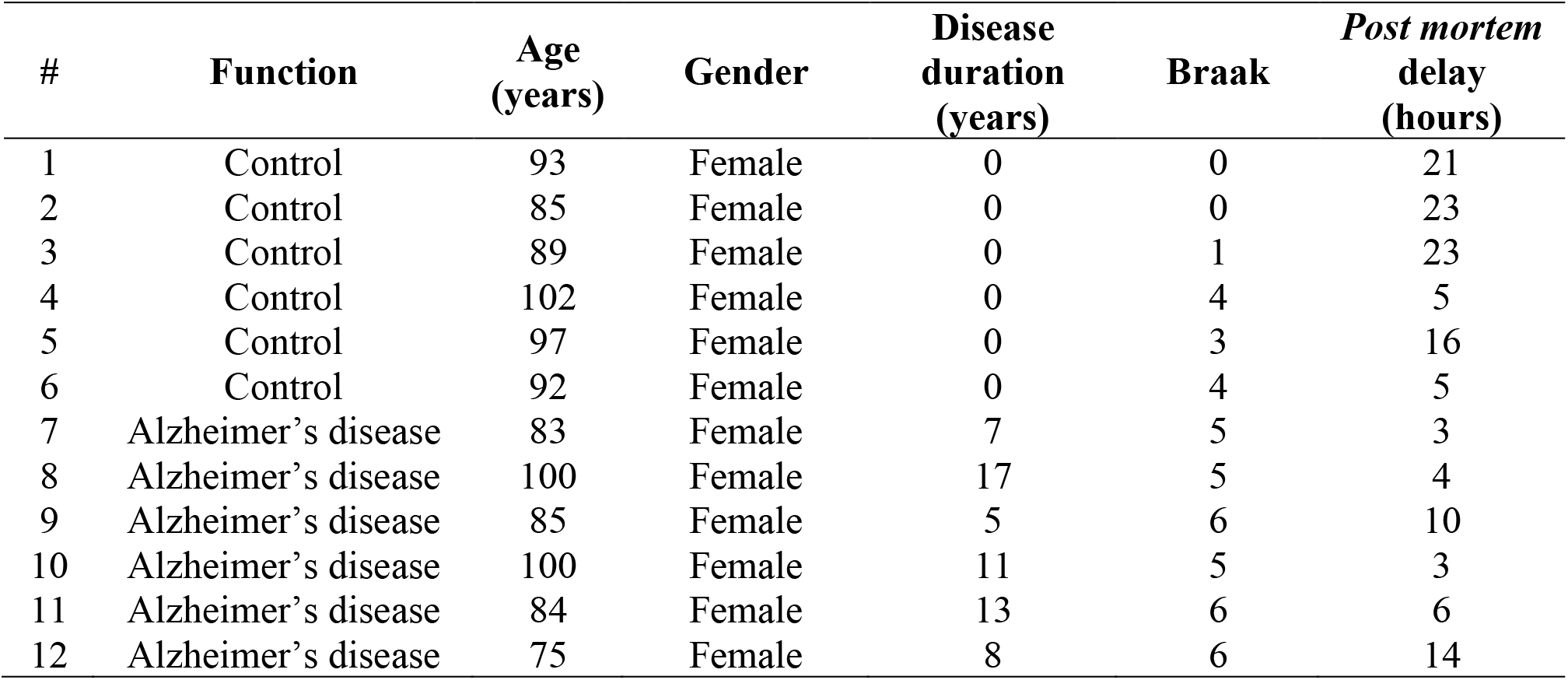
Human cases used for western blot analysis.

**Fig 4:**
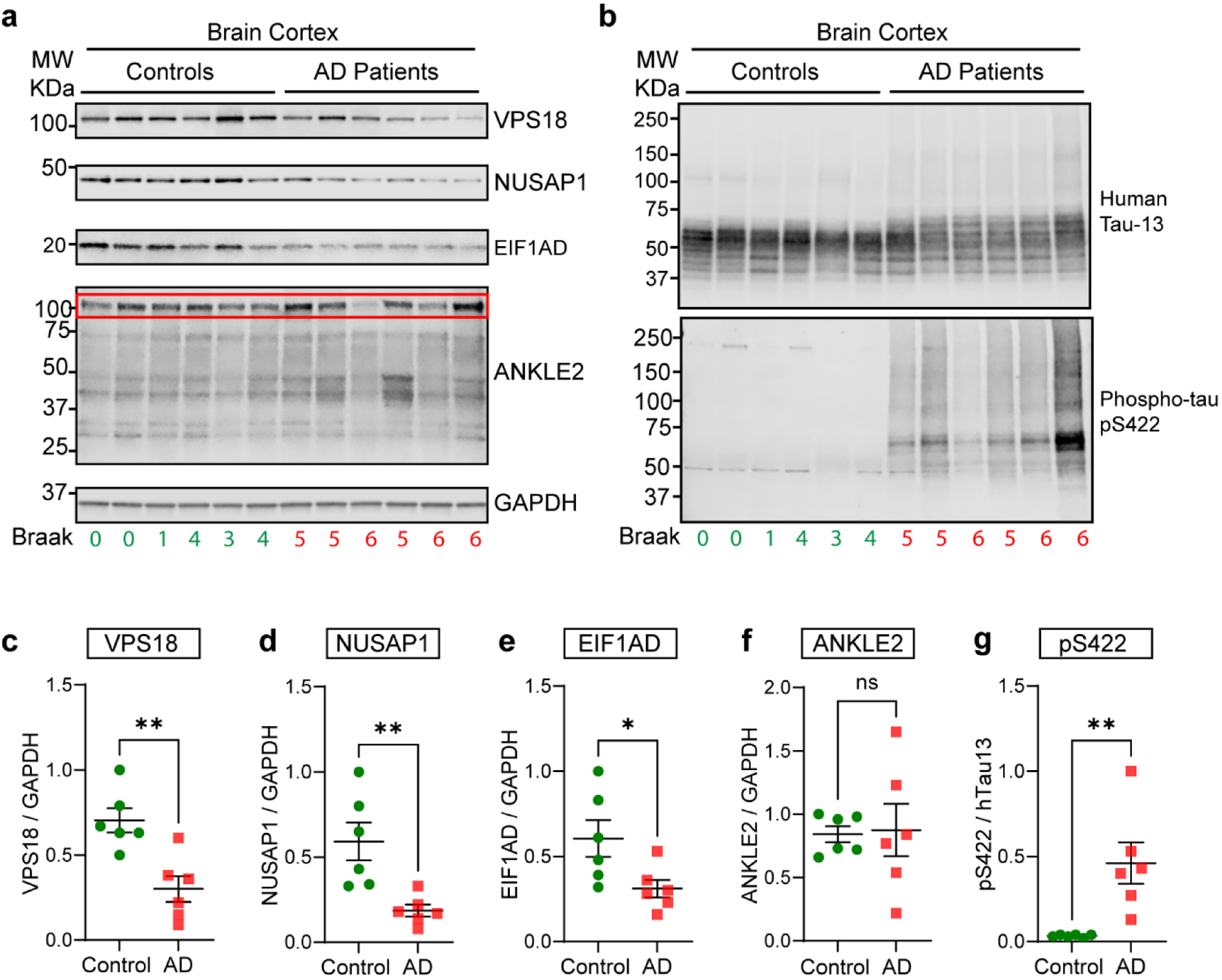
Quantitative western blot analysis of novel validated regulators in *post mortem* brain tissue from Alzheimer’ patients. (**a**) Western blot analysis of validated novel regulators using *post mortem* cortical brain samples from Alzheimer’s patients (AD). For the pathological data of the patients see Table 1. Antibodies against human VSP18, NUSAP1, EIF1AD and ANKLE2 were used. (**b**) Detection of human tau (Tau-13 antibody) and tau phosphorylated at Ser-422 (pS422 antibody) in the human brain samples. (**c-f**) Quantification of the protein levels in control and AD samples reveals that VPS18 (**c**), NUSAP1 (**d**) and EIF1AD (**e**) are significantly downregulated in AD patients. However, ANKLE2 levels (**f**) were not statistically different between both types of brain samples analyzed. Of note, the canonical isoform with a size of 104-117 kDa (red box outline) was the predominant isoform detected in human brains, and therefore, only this isoform was quantified. (**g**) Quantification of the ratio of pS422/hTau13 shows a substantial increase in tau phosphorylation at Ser-422 in AD samples.

## Discussion

Tau forms pathological aggregates in tauopathy but how these are generated is only incompletely understood. In the familial forms of the primary tauopathy FTLD-tau, the disease is caused by autosomal dominant mutations in the tau-encoding *MAPT* gene^25,26^, demonstrating that tau dysfunction by itself can lead to neurodegeneration and dementia. The vast majority of tauopathies, however, are sporadic^27^. Here, we identified and functionally validated ANKLE2, BANF1, VPS18, NUSAP1, and EIF1AD as cellular regulators whose knockdown predisposes cells to tau aggregation, and therefore, the products of these genes function as host restriction factors that control tau pathology. Importantly, given that our hits did not affect the uptake of tau seeds, these genes are presumed to be downstream of tau internalization, causing tau aggregation via cell-autonomous mechanisms. Furthermore, we showed that VPS18, NUSAP1, and EIF1AD are all downregulated in AD brain samples, supporting the notion that these regulators may be important in the development of tauopathy in the human brain.

Regarding risk genes, genome-wide association studies (GWAS) have been instrumental in identifying genes linked to an altered risk of developing tauopathies such as AD or FTLD-tau^28-30^. Paradoxically, when a total of 22 AD risk modifier genes revealed by GWAS studies were ablated in the cellular model of tau biosensor cells, the corresponding gene knockouts did neither affect the uptake of tau seeds nor the development of tau aggregates, indicating that these risk genes may not directly be involved in tau propagation^31^. In our study, we employed a functional approach using CRISPR-based genomics to identify genes that regulate seeded tau aggregation induced by vesicle-free and exosomal tau seeds, aiming at identifying disease-relevant genes and unveiling potential causal determinants in tauopathy.

Our analysis of the biological pathways in which the first 200 gene hits of our screen could be operating revealed that these genes are mostly involved in pathways of the late endosome, autophagosome, and tethering complexes coordinating endosome and lysosome fusion. These results support the notion that decreased autophagy or impaired late-endosome activity increases tau aggregation, perhaps due to an inability to degrade newly forming tau aggregates. For instance, VPS18 and CHMP6 (FDR<5%) were two validated positive hits with roles in vesicle-mediated protein trafficking to lysosomal compartments, and sorting of endosomal cargo proteins into endosomal multivesicular bodies. Similarly, CHMP6 and VPS13A from our top list (FDR<5%) were also identified in a previous report that used a CRISPRi library restricted to genes in the endosomal pathway^8^. Given that in our genome-wide screen (that was not limited to endosomal genes), these genes were also found, strengthens the confidence in the role of CHMP6 and VPS13A in regulating seeded tau aggregation. Interestingly, CHMP2B, a functional relative of CHMP6, has been linked to familial forms of FTLD-tau^32,33^. Similarly, VPS18 was found to be downregulated in human iPSC-derived cerebral organoid models derived from familial AD tissue^34^. VPS18 is a subunit of the mammalian homotypic fusion and vacuole protein sorting (HOPS) complex that regulates the fusion of endosomes and autophagosomes with lysosomes^35^. Thus, our study supports what has been suggested in previous investigations that autophagic and endolysosomal dysfunction are a driving force in the development of tauopathies and neurodegeneration more generally^7,30,36- 39^, highlighting the importance of endolysosomal and autophagic dysfunction as a risk factor in neurodegenerative diseases.

Intriguingly, gene silencing of ANKLE2 and BANF1 resulted in some of the most potent effects on tau aggregation, yet little is known about the role of these genes in tauopathies. ANKLE2 has been found downregulated in the transcriptomes from laser-captured CA1 neurons and microglia from AD brains, as well as in AD hippocampal homogenates^40^. However, we were unable to corroborate the downregulation of ANKLE2 by western blotting using cortical brain tissue from AD and control aging patients, possibly due to differences between hippocampal and cortical tissue. Although ANKLE2 was not downregulated in our study, this does not rule out a potential loss-of-function of ANKLE2 due to an altered post-translational modification such as phosphorylation or acetylation^41,42^, or a potential inactivation due to protein aggregation. We would speculate that one potential mechanism by which ANKLE2 may influence tau aggregation is through its binding to PP2A^43^, a known serine/threonine-protein phosphatase that directly regulates tau phosphorylation and physically binds to tau^44,45^, potentially acting together with PP2A to dephosphorylate tau and thereby reduce tau aggregation. Another mechanism by which ANKLE2 could affect tau pathology is through the regulation of integrity of the nuclear envelope, which can be affected by de/phosphorylation of BANF1 regulated by ANKLE2^43^, and reducing ANKLE2 levels disrupts and alters the nuclear envelope morphology^42,43^. Interestingly, anomalous invaginations of the nuclear envelope have been found in AD and FTLD-tau patients^46,47^, and tau accumulates close to these invaginations^47^, with evidence suggesting that liquid-liquid phase separation of tau at the nuclear envelope might be a possible initiating event in tauopathies^46^. Regarding BANF1, mutations in its gene can cause a human progeroid syndrome^48^, and it appears that BANF1 is crucial for restoring the capacity to repair oxidative lesions^49^. Unfortunately, we did not detect BANF1 in human brain samples by western blotting, possibly because BANF1 is a very small protein of only 10 kDa that could have been easily degraded during the *post mortem* interval of human tissue sampling or, alternatively, because the aging brain exhibits low levels of BANF1.

Like tau, NUSAP1 is a microtubule-stabilizing protein found in the cytoplasm and nucleolus. It localizes to the mitotic spindle in dividing cells and causes mitotic arrest when overexpressed^50^. Furthermore, a potential epistatic genetic interaction between the SNP rs16971798 of NUSAP1 and the presence of tau filaments has been reported^51^. The nucleolus is the site of ribosome biogenesis and rRNA processing^52^, and nucleolar dysfunction has been reported in AD patients as a potential link to the development of tauopathies^53-56^. Our study is the first to report that a loss of function of NUSAP1 can facilitate tau aggregation and that the protein is downregulated in AD brain samples, potentially leading to nucleolar dysfunction and thereby enhancing tauopathy.

Continuing with the theme of ribosomal dysfunction, we identified EIF1AD that is required for the final steps of 40S ribosomal subunit assembly as shown in human cells^57^. To our knowledge, there is again no report associating EIF1AD directly with tauopathies; however, EIF1AD has been associated with other brain diseases such as bipolar disorder^58^. We speculate that because of EIF1AD’s role in the formation of the ribosome, its downregulation could cause ribosomal dysfunction and propitiate anomalous protein aggregation, given that translational impairment has been associated with tauopathies^52,59^.

In summary, we have used CRISPR-based functional genomics to reveal novel genetic players that predispose cells to tau aggregation. Intriguingly, at least three gene products, VPS18, NUSAP1, and EIF1AD were found downregulated in AD brains, supporting their potential relevance in the emergence or propagation of tauopathy. Our data therefore raise the possibility that for some of the genes revealed on our screen mutations may be identified that are associated with human tauopathies. As an example of the predictive value of such studies, a link has been established between valosin-containing protein (VCP) and tau pathology in several cellular and animal models^60-62^ long before a specific VCP mutation was associated with human tauopathy^63^. Importantly, all genes validated from our screen had an impact on seeded tau aggregation initiated by both vesicle-free and exosomal tau seeds, implying that therapeutically targeting our protein hits would not render neurons exposed to the attack of either free-or membrane-bound tau but holistically cover both entry routes^7^, thus representing a more efficacious and unifying treatment strategy for tauopathies.

## Acknowledgments

This study was supported by the Estate of Dr Clem Jones AO, as well as a grant from the National Health and Medical Research Council of Australia [GNT1161133] to JCP and JG, and the State Government of Queensland (DSITI, Department of Science, Information Technology and Innovation). GB was supported by the Academy of Finland Research Fellowship [318434]. We thank Linda Cumner, Keisha Roffey, Tishila Palliyaguru, Trish Hitchcock, and the animal care team for animal maintenance, Virginia Nink, and Nadia de Jager at QBI FACS facility for flow cytometry assistance.

## Author Contributions

JCP and GB designed and performed the experiments, analyzed the data, interpreted the experiments, wrote the manuscript, and provided funding; YA, AF, GRH, and AB performed experiments, analyzed the data, and contributed to manuscript writing; MVR provided human brain samples with clinical data and Braak staging; JG contributed to experimental design and interpretation, edited the manuscript, and provided funding.

## Additional information

Supplementary information can be found in the online version available on *journal’s website*

## Materials & Correspondence

Further information and correspondence should be directed to. All unique resources generated in this study are available with a completed Materials Transfer Agreement.

## Methods

### Mouse strains and collection of brain tissue

Transgenic rTg4510 mice expressing human tau containing the P301L mutation that has been linked with familial frontotemporal dementia^22^ were used at 6-12 months of age for isolations of exosome and sarkosyl-insoluble tau from dissected brains. Animal experimentation was approved by the Animal Ethics Committee of the University of Queensland (approval number QBI/505/17/NHMRC and QBI/554/17/NHMRC).

### Plasmids and sgRNA cloning

The human Dolcetto CRISPR inhibition pooled library (Addgene #92385), and the plasmids pLX_311-KRAB-dCas9 (Addgene #96918), pLV hU6-sgRNA hUbC-dCas9-KRAB-T2a-Puro (Addgene #71236), psPAX2 (Addgene #12260), and pMD2.G (Addgene #12259) were a kind gift from John Doench, David Root, Charles Gersbach, and Didier Trono to Addgene. For phenotypic validation, each sgRNA hit (Supplementary Table S2) was individually cloned as annealed oligonucleotides into pLV hU6-sgRNA hUbC-dCas9-KRAB-T2a-Puro using Golden Gate cloning^64^ with FastDigest Esp3I (Thermo FD0454) and T4 DNA ligase (Thermo K1422). All oligonucleotides were purchased from IDT, and generated plasmids were corroborated by sequencing.

### Isolation of sarkosyl-insoluble tau from brains of tau transgenic mice

Biochemical isolation of sarkosyl-insoluble tau from brains of rTg4510 mice was performed as previously described^65^. Briefly, one brain was homogenized in 3 ml of ice-cold 1X RIPA buffer (150 mM NaCl, 50 mM Tris-HCl pH7.4, 0.5% (w/v) sodium deoxycholate, 1.0% (v/v) Nonidet P-40, 5 mM EDTA, 50 mM NaF, 200 mM NaVO4) containing 1x Complete protease inhibitor cocktail (Roche) using a drill-driven Teflon douncer. Homogenate was centrifuged at 20,000g for 20 min, the supernatant was mixed 1:1 with RIPA buffer with 2% sarkosyl (Sigma L9150) and incubated for 1 hour at room temperature with shaking. Insoluble tau was pelleted at 120,000g for 70 min, resuspended in 500 μl phosphate-buffered saline (PBS, Lonza 17-516Q) with protease inhibitors, and sonicated with three 10 sec pulses at 30% amplitude using a probe sonicator (Sonics Vibra-Cell). Sonicated sample was diluted with 3.5 ml PBS and concentrated to 500 μl by diafiltration using an Amicon® Ultra-4 Centrifugal Filter Unit with 30 KDa cutoff (Merck UFC803024) to remove traces of sarkosyl. Protein content was quantified with a BCA™ Protein Assay Kit (Thermo 23227).

### Isolation and purification of brain exosomes

Exosomes were isolated from the interstitial space of the mouse brain using a previously established protocol^10,13,15^. In brief, each brain was chopped, and the cells dissociated for 30 min at 37 °C with 0.2% collagenase type III (Worthington LS004182) in Hibernate-A medium (Thermo A1247501), followed by gentle pipetting with a 10 ml pipette. A series of differential 4 °C centrifugations at 300 g for 10 min, 2,000 g for 10 min, and 10,000 g for 30 min was then performed to discard the pellets containing cells, membranes, and nanodebris, respectively. The supernatant from the 10,000g centrifugation step was passed through a 0.22 μm syringe filter (Millipore Millex-GP) and ultracentrifuged at 120,000g for 70 min at 4 °C to pellet the exosomes. Pellets from five mouse brains per genotype were pooled, washed with PBS, and ultracentrifuged. This pooled preparation of exosome pellets was resuspended in 2 ml of 0.95 M sucrose in 20 mM HEPES (Thermo 15630080), after which a sucrose step gradient (six 2 ml steps: 2.0, 1.65, 1.3, 0.95, 0.6, and 0.25 M on top) was used to purify the exosomes by centrifugation at 200,000 g for 16 h at 4 °C. Finally, the sucrose-purified exosomes floating in the interphase between 0.95 M and 1.3M sucrose were recovered, washed with 5 ml PBS, ultracentrifuged again, and the exosome pellet resuspended in 120 μl PBS containing 1x Complete protease inhibitor cocktail (Roche) and 100 units/ml of Penicillin-Streptomycin (Thermo 15140122). Protein content was quantified with a BCA™ Protein Assay Kit using a 15 μl aliquot of exosomes in PBS, which was mixed with 15 μl of 1X RIPA buffer (150 mM NaCl, 50 mM Tris-HCl pH7.4, 0.5% (w/v) sodium deoxycholate, 1.0% (v/v) Nonidet P-40, 1% (w/v) SDS, 5 mM EDTA, 50 mM NaF) supplemented with protease inhibitors, and then homogenized in a water bath sonicator for 10 min.

### Fluorescent labeling of sarkosyl-insoluble tau and exosomes

Approximately 1 mg of sarkosyl-insoluble tau as 500 μl PBS solution at 2 mg/ml was fluorescently labeled on the N-terminus of protein aggregates using an Alexa-Fluor 647 Protein Labeling Kit (Thermo A20173) following the manufacturer’s instructions. For brain-derived exosomes, 600 μg protein equivalents of exosomes pelleted by ultracentrifugation were resuspended in 500 μl Diluent-C for membrane labeling (Sigma CGLDIL) and then mixed 1:1 with 500 μl Diluent-C containing 2 μl CellVue Claret Far-Red Fluorescent Membrane Linker (Sigma MINCLARET), labeling mixture that was incubated for 10 min at room temperature in the dark. Labeled exosomes were diluted with 3.5 ml PBS containing 1x Complete protease inhibitor cocktail (Roche) and concentrated to 300 ul by diafiltration using an Amicon® Ultra-4 Centrifugal Filter Unit with 30 kDa cutoff (Merck UFC803024) to remove the potentially unincorporated dye. Protein content of fluorescently labeled exosomes and sarkosyl tau was determined by BCA™ Protein Assay as described above.

### Culture of tau biosensor cells and HEK293 Lenti-X cells

The ‘tau biosensor cells’ are a monoclonal HEK293T cell line that stably expresses two fluorescently tagged forms of the microtubule-binding domain of tau bearing the P301S mutation, RD-CFP and RD-YFP, and was kindly provided by Dr. Marc Diamond^11^. BSKRAB cells were generated by transducing tau biosensor cells with lentiviral pLX_311-KRAB-dCas9 and used for CRISPRi genome-wide screening. BSKRAB-KD (individual knockdown) cells used in validation experiments were generated by transducing tau biosensor cells with lentiviral pLV hU6-sgRNA hUbC-dCas9-KRAB-T2a-Puro targeting each gene individually (Supplementary Table S2). Lenti-X 293T cells (Takara 632180) are a subclone of HEK293T which supports high levels of viral protein expression. All cells were grown in DMEM (Dulbecco’s modified Eagle’s medium, Thermo 11965092) supplemented with 100 units/ml of Penicillin-Streptomycin (Thermo 15140122), 2 mM GlutaMAX (Thermo 35050061), and 10% fetal bovine serum (FBS, Scientifix SFBS-FR).

### Library production

Human CRISPRi sgRNA library Dolcetto Set A^17^ (Addgene #92385) was transformed into electrocompetent Lucigen Endura™ *E. coli* (Lucigen 60242-2) using program EC1 on MicroPulser Electroporator (Bio-Rad 1652100) following the manufacturer’s instructions. The electroporated bacteria were plated onto 10 × 15 cm LB-agar dishes with 100 μg/ml ampicillin. After incubation for 16 h at 32°C, the bacteria were collected with a scraper in 5 ml PBS per dish, and plasmid DNA was extracted with the NucleoBond Xtra Midi kit (Macherey Nagel740410.50). The transformation efficiency was assessed by plating 1/10,000 of the reaction onto a 15 cm LB-agar plate with 100 μg/ml ampicillin.

### Production of lentiviral particles

Small scale production of active lentiviral particles was performed with 3^rd^ generation lentiviral transfer plasmids (500 ng each) mixed with 500 ng of a packaging DNA premix using psPAX2 and pMD2.G in a 2:1 ratio, which were transfected into Lenti-X 293T cells using TransIT-VirusGEN (Mirus MIR6700) according to the manufacturer’s instructions for a 12-well plate. Transfection mixture was added to Lenti-X 293T cells cultured in DMEM containing 10% FBS. Lentivirus-containing conditioned medium was collected after 60 hours, centrifuged at 1,000g for 5 min, and then filtered at 0.45 μm. Cells were transduced in DMEM medium supplemented with 10 mM HEPES and 8 μg/ml of Polybrene (Sigma H9268) immediately prior to the conditioned medium being added.

A larger-scale procedure was used for pooled CRISPRi library production in which Lenti-X 293T were seeded in 15 cm tissue culture dishes at a density of ∼ 10^5^ cells per cm^2^ overnight before transfection with the CRISPRi library (20μg/dish), packaging plasmids pMD2.G (5μg/dish), psPAX2 (25μg/dish), using the transfection reagent TransIT-LT1 (152μl/dish; Mirus MIR2300). The DNA mixture was suspended in 6 ml of DMEM (Thermo 11965092). The solution was incubated at room temperature for 20 min, during which time the growth medium was changed on the Lenti-X 293T cells. After this incubation, the transfection mixture was added dropwise to Lenti-X 293T cells, and the plates were incubated at 37 °C for 8 h. Transfection medium was then removed and replaced with DMEM +10% FBS supplemented with 0,5% BSA. Lentivirus-containing medium was collected 48h later, centrifuged at 3,000xg for 10 min at 4 °C, and the supernatant was aliquoted and stored at - 80 °C. The virus titer was determined by serial dilution in HEK293T cells followed by puromycin selection (1ug/ml) starting 48 h post infection. The number of puromycin-resistant cells was used as a measure of virus infectious units.

### Tau aggregation in tau biosensor cells transduced with a CRISPRi library

CRISPRi library cells were generated by transducing BSKRAB cells in four biological replicates at a low MOI (∼0.5) with the Dolcetto lentiviral library achieving an estimated representation of 1,000 cells per sgRNA per replicate. Transduced cells were selected with puromycin (1μg/ml) starting 48h post infection. For genome-wide screens, 13×10^6^ CRISPRi library cells were seeded in T175 flasks overnight. The following day, the medium was aspirated in cells at approximately 70% confluence, and then treated with 120 μg sarkosyl-insoluble tau (vesicle-free tau seeds) resuspended in 35 ml DMEM media supplemented with Pen/Strep and GlutaMAX as above but using 5% exosome-depleted fetal bovine serum (edFBS) prepared by centrifugation of FBS at 120,000 g for 18 h, followed by filter sterilization of the supernatant. CRISPRi library cells strongly developed tau inclusions by 48 h, at which time the cells were analyzed by FRET flow cytometry as described below.

### FRET flow cytometry

Tau aggregation between RD-CFP and RD-YFP was visualized and quantified by FRET flow cytometry as previously described^10,11,13^. In brief, CRISPRi library cells in T175 flasks used for the screens were harvested with 3 ml 0.25% trypsin-EDTA (Thermo 25200056) at 37°C for 5 min, mixed with 5 volumes culture medium, centrifuged at 300x for 5 min, supernatant aspirated, cell pellet washed with PBS, and then resuspended in ice-cold FACS buffer (PBS containing 30 mM HEPES, 0.5 mM EDTA and 0.2% BSA) prior to FRET flow cytometry using a FACSAria cell sorter (Becton Dickinson), where cells were excited by a 405 nm laser (Coherent Inc.) and the emitted fluorescence was captured with filters for 485/22 nm to detect CFP and 530/30 nm to detect FRET, gating the cells as outlined previously^10,13^. For genomic screens, all the FRET-positive and negative cells from T175 flasks were sorted and collected independently. For validation assays in 96-well plates, cells washed with PBS before being dissociated with 40 μl 0.25% trypsin-EDTA without phenol red (Thermo 15400054) and then mixed in the well with 160 μl DTI FACS buffer prepared with Defined Trypsin Inhibitor (Thermo R007100) supplemented with 30 mM HEPES, 0.5 mM EDTA and 0.2% BSA. FRET flow cytometry was performed as above, analyzing 40,000 cells per triplicate in each experiment. FRET data were quantified as the integrated FRET signal, calculated by multiplying the percentage of FRET-positive cells in the sample by their respective mean 530 nm fluorescence intensity generated by FRET.

### Validation of hits in individual knockdowns

For validation experiments, tau biosensor cells were transduced in 12-well plates with lentiviruses targeting each hit individually, by cloning each sgRNA into pLV hU6-sgRNA hUbC-dCas9-KRAB-T2a-Puro, and using non-targeting sgRNAs as a control (Supplementary Table S2). Each well contained an individual knockdown cell line (BSKRAB-KD), and these cells were split in DMEM culture medium with puromycin (1μg/ml) plus 10% edFBS after 72h transduction. Individual BSKRAB-KD cells from each 12-well were plated on 96-well plates in triplicates at a density of 20,000 cells per well overnight using 100 μl of puromycin-containing medium. On the next day, 50 μl of culture medium were removed and replaced with either 400 ng sarkosyl-insoluble tau or 3 μg protein equivalents of exosomes prepared in 50 μl fresh culture medium, incubating the treated BSKRAB-KD cells for further 72 h prior to FRET flow cytometry analysis as described above. Cells were topped up with 100 μl fresh medium at 48h to avoid acidification of the culture medium. In parallel, 600,000 BSKRAB-KD cells per individual knockdown were seeded in 6-wells and grown for 72 h, to prepare whole-cell lysates to corroborate knockdowns by western blots.

### Uptake quantification of exosomal and vesicle-free tau seeds

To measure levels of tau seed uptake, individual knockdown BSKRAB-KD cells were plated on 96-well plates at a density of 50,000 cells per well using 100 μl DMEM medium plus edFBS. 48 hours later, 50 μl of culture medium were removed and replaced with either 400 ng sarkosyl-insoluble tau labeled with Alexa-Fluor 647 or 2 μg protein equivalents of exosomes labeled with far-red CellVue Claret prepared in 50 μl fresh culture medium with edFBS. These cells were incubated at 37 °C for 60 min, then the medium was aspirated, the cells washed with PBS, dissociated with 40 μl 0.25% trypsin-EDTA without phenol red (Thermo 15400054) at 37°C for 10 min, and finally mixed with ice-cold 160 μl Defined Trypsin Inhibitor (Thermo R007100). To remove potential traces of non-internalized labeled tau seeds, trypsin-dissociated cells on 96-well plates were centrifuged at 1,000g for 5 min at 4°C using a swinging-bucket rotor for microplates (Beckman S6096) in an Allegra X-30R centrifuge (Beckman). The supernatant was aspirated after centrifugation and cell pellets resuspended in 200 μl ice-cold DTI FACS buffer (formulation above) prior to flow cytometry using an BD® LSR II. Tau uptake was quantified by measuring the mean fluorescence intensity in far-red positive cells, analyzing 40,000 cells per triplicate in each experiment.

### Genomic DNA preparation and next-generation sequencing

Genomic DNA from FRET-positive and negative cells were isolated using commercial kits (QIAGEN 13323 and 13343). We amplified sgRNA cassettes from gDNA (2.5μg) using. OneTaq® DNA Polymerase (NEB #M0480) and LG.Lib.ampl1.F and LG.Lib.ampl1.R primers in a 50 μl reaction. Then, Illumina sequencing primer binding sites were added by PCR amplification of sgRNA amplicons with primer mix LG.LibAmpl.WSstag.mix and LG.gRNA.Ampl.NGS.R. Lastly, Illumina indices and adapters for sample multiplexing were added by PCR amplification with Illumina_indX_F and Illumina_indX_R primers. The last two PCR rounds were performed using NEBNext® Ultra™ II Q5® Master Mix (NEB #M0544). Samples were purified using AMPure XP beads (Beckman Coulter #A63880). The library was sequenced with coverage of 200 reads per sgRNA using the PE100 protocol on NextSeq500 Illumina sequencer (with 10% PhiX spike-in). Samples were demultiplexed and spacers were counted using the count_spacers.py script^66^. Positively selected genes were identified using the MAGeCK tool^24^. The over-representation and gene set enrichment analysis for GO-BP (biological process) and GO-CC (cellular component) terms were performed using clusterProfiler R package^67^. Primer sequences and detailed reaction setups are listed in Supplementary Table S3.

### Bioinformatic Analysis

Samples were demultiplexed using Illumina bcl2fastq to generate FASTQ files. Individual sgRNA counts were extracted using the count_spacers.py script^66^. Positively selected genes were identified using the MAGeCK tool^24^ and DESeq2(Wald)^68^ using simplified routines provided by DEBRA R package^69^. The over-representation and gene set enrichment analysis for GO-BP (biological process) and GO-CC (cellular component) terms were performed with clusterProfiler R package^67^ using the first 200 genes with the following parameters pAdjustMethod = “BH”, pvalueCutoff = 0.25, qvalueCutoff = 0.25.

### Brain and cell lysates

HEK293T cells grown in 6-well plates were used to prepare whole-cell lysates using a pellet of trypsin-dissociated cells that was homogenized in 200 μl RIPA buffer (150 mM NaCl, 50 mM Tris-HCl pH7.4, 0.5% (w/v) sodium deoxycholate, 1.0% (v/v) Nonidet P-40, 1% (w/v) SDS, 5 mM EDTA, 50 mM NaF) with protease inhibitors using a probe sonicator (Sonics Vibra-Cell) for 20 sec at 30% amplitude. Sonicated lysates were left solubilizing on ice for 1 h, then centrifuged at 20,000g for 20 min, using the supernatant for protein quantification with a BCA™ Protein Assay. Similarly, human brain lysates from *post mortem* AD patients were prepared with 25 mg of cortical brain tissue homogenized in 500 μl RIPA buffer supplemented with protease and phosphatase inhibitors, disrupting the tissue with 20 strokes of a drill-driven Teflon Dounce homogenizer, solubilizing the lysates on ice for 30 min and then centrifuged at 20,000g for 20 min, taking the supernatant for protein quantification.

### Western blot analysis

Criterion TGX 4-15% (Bio-Rad 5671084) and Mini-Protean TGX precast gels (Bio-Rad 4561083) were used to separate 20-40 μg of total protein from lysates, which were then transferred onto Immuno-Blot low fluorescence PVDF membranes (Bio-Rad 1704275) using the Trans-Blot Turbo transfer system (Bio-Rad). Membranes were blocked in Odyssey Blocking Buffer (Li-Cor) for 1 h at room temperature (RT) and then incubated overnight at 4°C with primary antibodies prepared in Odyssey Blocking Buffer with 0.01% Tween-20. Membranes were washed with Tris-buffered saline/0.1% Tween-20 (TBST) three times for 10 min at RT. Then, IRDye secondary antibodies (Li-Cor) were added diluted 1:10,000 in a 1:1 mixture of Odyssey Blocking Buffer with TBST for 1 h at RT. Finally, membranes were again washed three times in TBST, and the fluorescence signals were recorded using an Odyssey FC imaging system (Li-Cor). Analysis and protein quantification were performed using Image Studio software (Li-Cor). The following antibodies were used: Anti-human ANKLE2 rabbit polyclonal (1:1,000; Thermo A302-965A), anti-human EIF1AD rabbit polyclonal (1:1,000; Thermo 20528-1-AP), anti-BANF1/BAF rabbit monoclonal (1:1,000; Abcam ab129184), anti-human NUSAP1 rabbit polyclonal (1:500; Abcam ab137230), anti-human VPS18 rabbit monoclonal (1:1,000; Abcam ab178689), anti-human Tau-13 mouse monoclonal (1:1,000; Covance MMS-520R), anti Tau phospho Ser422 rabbit polyclonal (1:1,000; GeneTex GTX86147), and the normalizers anti-GAPDH mouse monoclonal (1:2,000; Millipore MAB374) and anti-GAPDH rabbit polyclonal (1:2,000; Proteintech 10494-1-AP).

### Statistical analysis

To determine the statistical significance of differences in quantification levels in validation experiments, *p*-values either were determined from a two-tailed unpaired *t*-test with Welch’s correction or from one-way ANOVA analysis with a 95% confidence interval and Dunnett’s test to correct for multiple comparisons, calculated with GraphPad Prism v9.3 for Windows (GraphPad Software Inc).

