## Supplementary material for "Genome-wide CRISPRi screening reveals regulators of Alzheimer’s tau pathology shared between exosomal and vesicle-free tau seeds": Supp FigS1, Supp Tables S1-S3

**Supplementary FigS1**

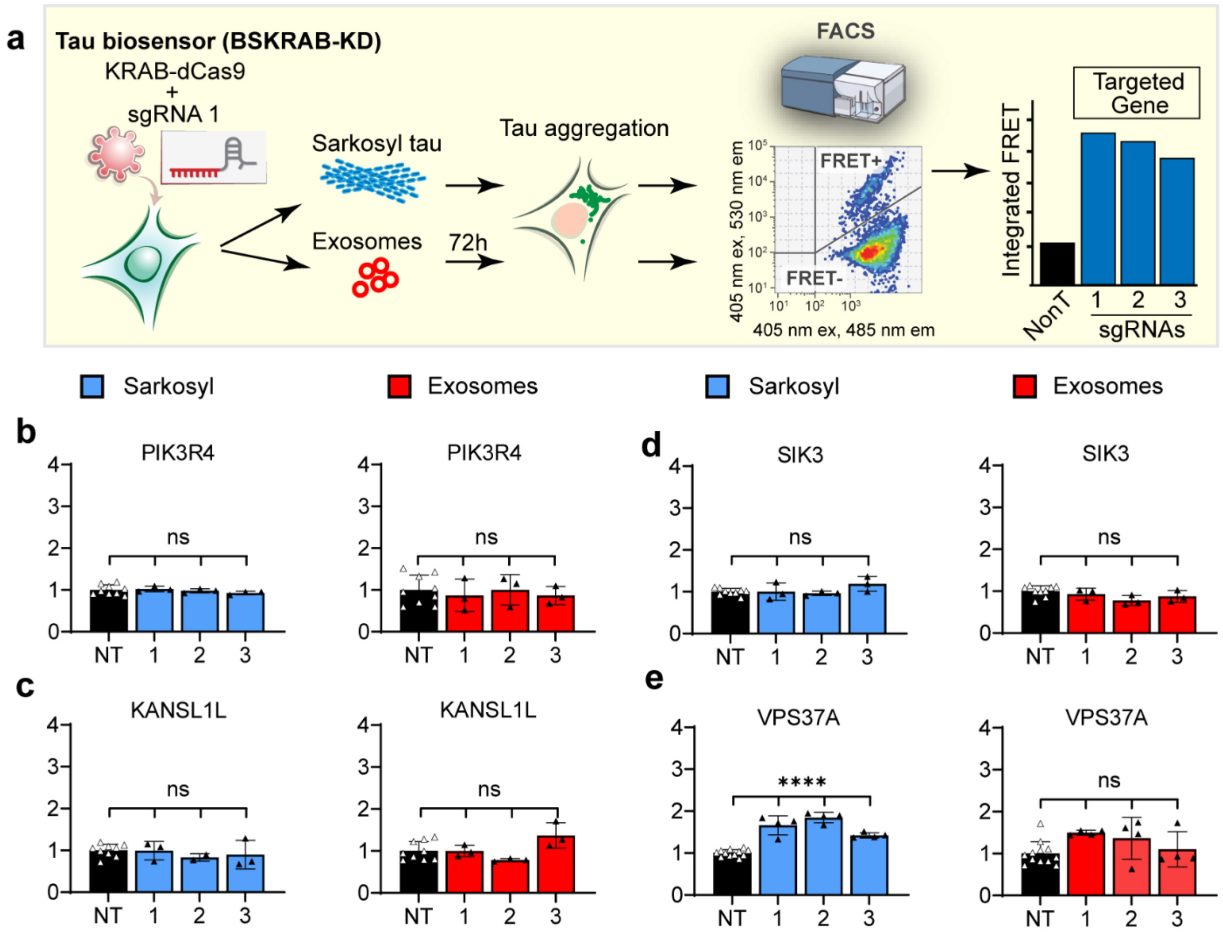


**Fig S1: Additional bioinformatic hits that were functionally validated using individual CRISPRi knockdowns. (a)** Workflow for functional validation. Individual sgRNAs were used to silence the targeted gene in combination with KRAB-dCas9 in tau biosensor cells (BSKRAB-KD), then treated with either sarkosyl-insoluble tau or exosomal tau seeds for 72 hours, followed by detection and quantification of tau aggregation using FRET flow cytometry. **(b-e)** Integrated FRET intensities represent levels of tau aggregation under knockdown conditions for the different gene hits. The control black column (NT) is the average obtained with three independent non-targeting sgRNAs (n=3) assessed in triplicates. Control cells were compared with knockdown cells targeted individually (1, 2, and 3). Individual silencing of PIK3R4, SIK3, and KANSL1L did not result in a significant increase in tau aggregation. Knockdown of VPS37A produced a weak potentiation of tau aggregation, in particular for exosomal tau. Error bars represent SEM for n=3; ****$p<0.0001$; ns, not significant.

**Supplementary Table S1:** List of top hits with %FDR<25. Hits were organized according FDR. LogFC, and RRA are also shown. Gray shadow highlights genes that were selected for functional validation experiments. *Genes previously identified by Chen and colleagues (*J Biol Chem* 2019).

| # Ranking | Gene | %FDR | logFC | RRA |
|---|---|---|---|---|
| 1 | BANF1 | **0.00** | 3.266637537 | 1.01498E-05 |
| 2 | EIF1AD | **0.00** | 1.934508458 | 1.47472E-08 |
| 3 | NUSAP1 | **0.00** | 2.031604495 | 7.46533E-08 |
| 4 | ATL2 | **0.00** | 1.773063362 | 1.4612E-05 |
| 5 | ANKLE2 | **0.00** | 1.278130789 | 2.07356E-07 |
| 6 | FOS | **0.36** | 1.691459075 | 0.000351751 |
| 7 | VPS18 | **0.36** | 1.352141849 | 5.07635E-05 |
| 8 | C1QA | **0.36** | 2.426433473 | 0.005616097 |
| 9 | CKAP2L | **0.78** | 1.519552533 | 0.000368053 |
| 10 | PIK3R4 | **0.78** | 0.789122106 | 2.84006E-07 |
| 11 | BAAT | **0.78** | 3.031695148 | 0.021070745 |
| 12 | SPNS1 | **1.10** | 2.251754864 | 0.006244625 |
| 13 | PRPS1L1 | **1.10** | 1.447539509 | 0.000420625 |
| 14 | CHMP6* | **1.23** | 1.227152175 | 0.000143875 |
| 15 | KANSL1L | **1.57** | 1.860221923 | 0.003519089 |
| 16 | FXR2 | **1.57** | 1.298024251 | 0.00031548 |
| 17 | ANAPC2 | **1.57** | 1.506391407 | 0.000998791 |
| 18 | VPS13A* | **1.57** | 1.143422967 | 0.000116332 |
| 19 | VARS | **1.57** | 1.514069683 | 0.00126152 |
| 20 | TTF2 | **1.57** | 0.989256494 | 3.7522E-05 |
| 21 | TBC1D16 | **2.21** | 1.491779155 | 0.001366599 |
| 22 | LCOR | **2.21** | 1.622711278 | 0.002364479 |
| 23 | SIK3 | **4.61** | 1.900795461 | 0.006582789 |
| 24 | VPS37A | **8.59** | 1.261569991 | 0.000736016 |
| 25 | HNF1A | **9.47** | 1.098094694 | 0.00029412 |
| 26 | ACAA2 | **9.47** | 1.196523181 | 0.000630893 |
| 27 | TARBP1 | **11.66** | 1.260472032 | 0.001156434 |
| 28 | IQCB1 | **11.66** | 0.992898619 | 0.000199406 |
| 29 | NSRP1 | **13.46** | 1.275077258 | 0.001462328 |
| 30 | CIR1 | **13.46** | 1.247949149 | 0.001398973 |
| 31 | SNRNP40 | **15.25** | 0.97273383 | 0.000242998 |
| 32 | EPHB6 | **15.25** | 0.881470148 | 0.000121608 |
| 33 | C12orf74 | **15.25** | 1.271398863 | 0.001996916 |
| 34 | UGT1A4 | **15.25** | 1.003761708 | 0.000406877 |
| 35 | DHX15 | **15.25** | 1.058384713 | 0.000634876 |
| 36 | ITSN1 | **15.25** | 1.55963337 | 0.007230958 |
| 37 | RMDN3 | **15.25** | 0.847998378 | 0.000119616 |
| 38 | CCNI | **15.25** | 1.016171052 | 0.000552394 |
| 39 | RABL3 | **15.25** | 0.819676089 | 9.40336E-05 |
| 40 | GLCE | **15.25** | 0.954770535 | 0.0003647 |
| 41 | LRRC49 | **16.55** | 0.827478125 | 0.000137699 |
| 42 | MUM1 | **21.07** | 1.089692807 | 0.001410185 |
| 43 | IL33 | **22.58** | 1.20655341 | 0.002856526 |
| 44 | VPS41 | **22.58** | 0.713229649 | 5.18977E-05 |
| 45 | MET | **23.61** | 1.508942865 | 0.009782958 |
| 46 | ETFDH | **24.59** | 1.022722103 | 0.001208978 |

**Supplementary Table S2**: Oligonucleotides encoding sgRNAs for individual knockdowns of hits. Annealed oligos were cloned into the BsmBI restriction sites of pLV hU6-sgRNA hUbC-dCas9-KRAB-T2a-Puro (Addgene #71236).

| **Gene targeted_sgRNA#** | **Sense oligo** | **Antisense oligo** |
|---|---|---|
| NON-TARGET_ sgRNA1 | CACCGAAAAAGCTTCCGCCTGATGG | AAACCCATCAGGCGGAAGCTTTTTC |
| NON-TARGET_ sgRNA2 | CACCGAAAATCGATGGGCTGAATCT | AAACAGATTCAGCCCATCGATTTTC |
| NON-TARGET_ sgRNA3 | CACCGAAAATTATCGGAAACGGTAG | AAACCTACCGTTTCCGATAATTTTC |
| ANKLE2_sgRNA1 | CACCGCCCGGGCGGCGGCGATGCTG | AAACCAGCATCGCCGCCGCCCGGGC |
| ANKLE2_sgRNA2 | CACCGGCTGGCGGCGGCCGAGTGGG | AAACCCCACTCGGCCGCCGCCAGCC |
| ANKLE2_sgRNA3 | CACCGCCACAGCATCGCCGCCGCCC | AAACGGGCGGCGGCGATGCTGTGGC |
| BANF1_sgRNA1 | CACCGGTTACGGGAACTGAAGTTGC | AAACGCAACTTCAGTTCCCGTAACC |
| BANF1_sgRNA2 | CACCGAGGTATCCGCAGGAGCGGCC | AAACGGCCGCTCCTGCGGATACCTC |
| BANF1_sgRNA3 | CACCGCCGCAGGAGCGGCCGGGTGG | AAACCCACCCGGCCGCTCCTGCGGC |
| EIF1AD_sgRNA1 | CACCGTCAGCGTCCAGCCTAGACGG | AAACCCGTCTAGGCTGGACGCTGAC |
| EIF1AD_sgRNA2 | CACCGGAGGGCCAGAGACTAAGTGT | AAACACACTTAGTCTCTGGCCCTCC |
| EIF1AD_sgRNA3 | CACCGGCAGCGTTCCTGGGAGCCGA | AAACTCGGCTCCCAGGAACGCTGCC |
| NUSAP1_sgRNA1 | CACCGTCCCGGCGATACTCGGAAGA | AAACTCTTCCGAGTATCGCCGGGAC |
| NUSAP1_sgRNA2 | CACCGATCATCGCGATTCGAAATCC | AAACGGATTTCGAATCGCGATGATC |
| NUSAP1_sgRNA3 | CACCGATCCATCTTCCGAGTATCGC | AAACGCGATACTCGGAAGATGGATC |
| CHMP6_sgRNA1 | CACCGTGACGCGGCTCTGCTTCTTG | AAACCAAGAAGCAGAGCCGCGTCAC |
| CHMP6_sgRNA2 | CACCGGCCGAACAGGTTACCCATGG | AAACCCATGGGTAACCTGTTCGGCC |
| CHMP6_sgRNA3 | CACCGGAGCGGGAGGACCCGAGCTA | AAACTAGCTCGGGTCCTCCCGCTCC |
| PIK3R4_sgRNA1 | CACCGCCAGCAAACGCCGAACTCCC | AAACGGGAGTTCGGCGTTTGCTGGC |
| PIK3R4_sgRNA2 | CACCGCGAACTCCCGGGAAAGCAAC | AAACGTTGCTTTCCCGGGAGTTCGC |
| PIK3R4_sgRNA3 | CACCGGCGGTCTGCACTTCTCTCCC | AAACGGGAGAGAAGTGCAGACCGCC |
| VPS18_sgRNA1 | CACCGAATCACAGGCTCCCTTCAGC | AAACGCTGAAGGGAGCCTGTGATTC |
| VPS18_sgRNA2 | CACCGAGGTTGGGATCACCTGGCAC | AAACGTGCCAGGTGATCCCAACCTC |
| VPS18_sgRNA3 | CACCGGGGGCGAGGTTGGGATCACC | AAACGGTGATCCCAACCTCGCCCCC |
| VPS37A_sgRNA1 | CACCGGCTGGCCGGTTTGGGCGTCT | AAACAGACGCCCAAACCGGCCAGCC |
| VPS37A_sgRNA2 | CACCGTTTGGGCGTCTGGGCCGTGA | AAACTCACGGCCCAGACGCCCAAAC |
| VPS37A_ sgRNA3 | CACCGCACGGCCCAGACGCCCAAAC | AAACGTTTGGGCGTCTGGGCCGTGC |
| KANSL1L_sgRNA1 | CACCGGCGGGCGGACTGTTAAACCG | AAACCGGTTTAACAGTCCGCCCGCC |
| KANSL1L_sgRNA2 | CACCGCCGCTGCCCGCAGCTCCGTG | AAACCACGGAGCTGCGGGCAGCGGC |
| KANSL1L_sgRNA3 | CACCGCTGTTAAACCGCGGCGGCGG | AAACCCGCCGCCGCGGTTTAACAGC |
| SIK3_sgRNA1 | CACCGGGCTCCCCCAGTCCCGGCCC | AAACGGGCCGGGACTGGGGGAGCCC |
| SIK3_sgRNA2 | CACCGCGGAGCTGGCGGGGCTGCCG | AAACCGGCAGCCCCGCCAGCTCCGC |
| SIK3_sgRNA3 | CACCGGCGGCGGCGAGCGGAGCTGG | AAACCCAGCTCCGCTCGCCGCCGCC |

**Supplementary Tables S3:** Primer sequences and detailed reaction setups for next-generation sequencing.

**Primers**

| | |
|---|---|
| LG.Lib.ampl1.F | TTGTGGAAAGGACGAAACACCG |
| LG.Lib.ampl1.R | TCTACTATTCTTTCCCCTGCACTGT |
| LG.gRNA.Ampl.NGS.R | GTGACTGGAGTTCAGACGTGTGCTCTTCCGATCTNNNNNNNNNNNNTCTACTATTCTTTCCCCTGCACTGT |
| Illumina_indX_F | AATGATACGGCGACCACCGAGATCTACACNNNNNNNNACACTCTTTCCCTACACGACGCTCTTCCGATC*T |
| Illumina_indX_R | CAAGCAGAAGACGGCATACGAGATNNNNNNNNGTGACTGGAGTTCAGACGTGTGCTCTTCCGATC*T |

**WS Stager Mix**

| | |
|---|---|
| LG.LibAmpl.WSstag.3 | ACACTCTTTCCCTACACGACGCTCTTCCGATCTSWSWSWSWSWSWTTGTGGAAAGGACGAAACACCG |
| LG.LibAmpl.WSstag.5 | ACACTCTTTCCCTACACGACGCTCTTCCGATCTSWSWSWSWSWSWSWTTGTGGAAAGGACGAAACACCG |
| LG.LibAmpl.WSstag.7 | ACACTCTTTCCCTACACGACGCTCTTCCGATCTSWSWSWSWSWSWSWSWTTGTGGAAAGGACGAAACACCG |
| LG.LibAmpl.WSstag.9 | ACACTCTTTCCCTACACGACGCTCTTCCGATCTSWSWSWSWSWSWSWSWSWTTGTGGAAAGGACGAAACACCG |
| LG.LibAmpl.WSstag.2.1 | ACACTCTTTCCCTACACGACGCTCTTCCGATCTWSWSWSWSWSWTTGTGGAAAGGACGAAACACCG |
| LG.LibAmpl.WSstag.4.1 | ACACTCTTTCCCTACACGACGCTCTTCCGATCTWSWSWSWSWSWSWTTGTGGAAAGGACGAAACACCG |
| LG.LibAmpl.WSstag.6.1 | ACACTCTTTCCCTACACGACGCTCTTCCGATCTWSWSWSWSWSWSWSWTTGTGGAAAGGACGAAACACCG |
| LG.LibAmpl.WSstag.8.1 | ACACTCTTTCCCTACACGACGCTCTTCCGATCTWSWSWSWSWSWSWSWSWSTTGTGGAAAGGACGAAACACCG |

**Reaction 1**

| Reagent | Volume |
|---|---|
| OneTaq Buffer | 10 ul |
| dNTP | 1ul |
| LG.Lib.ampl1.F | 5 ul |
| LG.Lib.ampl1.R | 5 ul |
| Genomic DNA (2ug) | - |
| Betaine (5M) | 10ul |
| OneTaq polymerase | 0,25 |
| $H_2O$ | up to 50ul |

**Cycling conditions (Reaction 1)**

| Temperature | Time | Cycles |
|---|---|---|
| 94ºC | 10 min | 1 |
| 94ºC | 1 min | 25 |
| 55ºC | 1 min | |
| 68ºC | 1 min | |
| 68ºC | 2 min | 1 |
| 4ºC | hold | 1 |

**Reaction 2**

| Reagent | Volume |
|---|---|
| NEBNext® Ultra™ II | 10ul |
| WS Stager Mix | 1ul |
| LG.gRNA.Ampl.NGS.R | 1ul |
| Reaction 1 | 1ul |
| H2O | 7ul |

**Cycling conditions (Reaction 2)**

| Temperature | Time | Cycles |
|---|---|---|
| 98ºC | 2 min | 1 |
| 98ºC | 30 sec | 8 |
| 67ºC | 40 sec | |
| 72ºC | 40 sec | |
| 72ºC | 1 min | 1 |
| 4ºC | hold | 1 |

**Reaction 3**

| Reagent | Volume |
|---|---|
| NEBNext® Ultra™ II | 10ul |
| SB70X | 1ul |
| SB50X | 1ul |
| Reaction 2 | 1ul |
| H2O | 7ul |

**Cycling conditions (Reaction 3)**

| Temperature | Time | Cycles |
|---|---|---|
| 98ºC | 2 min | 1 |
| 98ºC | 30 sec | 8 |
| 72ºC | 1 min | |
| 72ºC | 1 min | 1 |
| 4ºC | hold | 1 |
